# Genomic factors contributing to the resilience of *Salmonella enterica* on ready-to-eat muskmelon

**DOI:** 10.1101/2025.06.02.657348

**Authors:** Irene Esteban-Cuesta, Laura Führer, Steffen Porwollik, Weiping Chu, Steven R. Fiddaman, Irmak Sah, Michael McClelland, Claudia Guldimann

## Abstract

*Salmonella* outbreaks have repeatedly been associated with muskmelons. To identify genes under selection in *S. enterica* growing in this food matrix, barcoded transposon mutant libraries in three *S. enterica* serovars *-* Typhimurium, Enteritidis, and Newport - were screened for survival and growth on muskmelon. Applying stringent thresholds, a total of 26 genes in Typhimurium, 34 in Enteritidis, and 50 in Newport were found to significantly influence fitness during muskmelon interaction, with many of these being temperature dependent. Genes whose disruption affected fitness across all three serovars were enriched for functions related to RNA degradation and ribosome biogenesis. Targeted competition assays confirmed the contribution of selected genes, revealing nutrient-dependent phenotypes for most mutants. Remarkably, the polyribonucleotide nucleotidyltransferase gene, *pnp,* and the D-3-phosphoglycerate dehydrogenase gene, *serA*, conferred a selective advantage when growing in muskmelon but not under nutrition-rich control conditions. In contrast, the nitrogen regulation response regulator GlnG provided a muskmelon-specific fitness disadvantage. This study provides novel insights into genome-wide adaptation mechanisms of multiple *Salmonella* serovars to growth on muskmelons, revealing both shared and serovar-specific determinants while illustrating the dynamic genetic responses of *S. enterica* throughout the interaction period.

## 1. Introduction

Non-typhoidal salmonellosis is a zoonotic disease that has a high impact on the economy and human and animal health. It is among the most relevant foodborne pathogens in the European Union (EU) and worldwide (EFSA and ECDC, 2024).

Ready-to-eat (RTE) melons pose a high infection risk to consumers since pathogen inactivation measures are rarely applied before consumption, and handling may enable the spread of surface contaminants to the inside of the fruit. *S. enterica* foodborne outbreaks associated with the consumption of muskmelons have been reported since 1990 (Ries, 1990; Bowen et al., 2006), with recent cases in the EU (2023)(McGeoch et al., 2024) and the United States (2024)(CDC, 2024). Especially, netted rind melon varieties offer a favorable niche for bacterial attachment and persistence (Fu et al., 2020).

To establish infection, *S. enterica* must be able to survive and grow on the food matrix. Quantitative data for the growth of *S. enterica* on muskmelons show that *S. enterica* can grow on muskmelons at 8°C and sustain its population at temperatures as low as 4°C (Ukuku and Sapers, 2007; BfR, 2013; Huang et al., 2015).

*S. enterica* responses to food-related stresses have been studied in the past (Spector and Kenyon, 2012; Pradhan and Devi Negi, 2019) and previous studies revealed fitness effects of gene disruptions vary significantly by food matrix, identifying key survival mechanisms on tomatoes (de Moraes et al., 2017; de Moraes et al., 2018), alfalfa sprouts (Holden et al., 2024), and low-water-activity foods, such as pistachios (Jayeola et al., 2020), black peppercorns, almonds, and hazelnuts (Li et al., 2020). Transposon insertion sequencing (TIS) showed matrix-specific effects for *Salmonella* colonization and persistence by the inactivation of genes linked to the alternative sigma factor σ^S^ (RpoS), DNA recombination, the biogenesis of lipopolysaccharides (LPS)(Jayeola et al., 2020), glycolysis, amino acid and LPS production, fatty acid degradation, and purine and pyrimidine biosynthesis pathways (de Moraes et al., 2018).

Based on the existing data for other food matrices, we hypothesized that distinct genetic determinants may exist for *S*. *enterica* fitness on muskmelon. In this study, we used barcoded transposon libraries of *S.* Typhimurium 14028s, *S.* Enteritidis PT4 strain P125109, and *S*. Newport C4.2, to identify genome-wide fitness effects of specific genes and the corresponding metabolic pathways involved in the interaction of these *S. enterica* serovars with RTE muskmelon. This interaction was examined under conditions relevant to retail and consumer handling, including storage at room temperature and during abusive cold storage at 8°C, a temperature that may be encountered both in open cooling displays at supermarkets and in household refrigerators (Laguerre et al., 2002; Evans and Redmond, 2015; Monge Brenes et al., 2020).

## 2. Material and Methods

### 2.1 Bacterial strains and culture media

*S. enterica* wild type (WT) strains used in this study and their sources are listed in **Table 1**. All strains were preserved at −80°C in Luria-Bertani broth (LB; Carl Roth GmbH & Co. KG, Karlsruhe, Germany) with 20% glycerol (Th. Geyer GmbH & Co. KG, Renningen, Germany). Strains were grown in LB broth or 1.5% LB agar (Oxoid Deutschland GmbH, Wesel, Germany). Where necessary, culture media were supplemented with 60 μg/ml kanamycin (LB^kan^; Carl Roth), 20 μg/ml chloramphenicol (LB^cm^, Carl Roth) or 15 μg/ml tetracycline (LB^tet^; Carl Roth).

**Table 1.** *Salmonella enterica* wild type strains used for the barcoded transposon libraries.

| Strain | Abbreviation | Source |
| --- | --- | --- |
| *Salmonella* Typhimurium strain<br>ATCC 14028 | STM | Isolated from 4-week-old chickens in 1960(Jarvik et al., 2010)<br>NCBI accession no. CP001363.1/ CP001362.2 |
| *Salmonella* Enteritidis PT4 strain<br>P125109 | SEN | Clinical strain associated with poultry outbreak in the United Kingdom(Toro et al., 2016)<br>NCBI accession no. CP063700.1/ CP063701.1 |
| *Salmonella* Newport C4.2 | SNP | Isolated from a tomato field in Virginia, United States(de Moraes et al., 2018)<br>NCBI accession no. GCA_006821125.1 |

### 2.2 Library construction

The construction of the barcoded Tn*5*-based mutant libraries has been previously described (de Moraes et al., 2017; de Moraes et al., 2018). Briefly, a library of Tn*5* insertion mutants was constructed for each strain with a mini-Tn*5* derivative into which an N_18_ random barcode was inserted via PCR. This construct was randomly introduced into the genome with the EZ-Tn*5* <T7/KAN-2> promoter insertion kit (Epicentre Biotechnologies, Madison, WI, United States). Transformed cells were recovered on LB^kan^ agar after overnight culture at 37°C.

The methods for mapping the barcoded transposons to specific locations in the genome have also been previously described (de Moraes et al., 2017). The *S.* Typhimurium ATCC 14028 transposon mutant library consisted of about 230,000 insertion mutants (de Moraes et al., 2017), the *S.* Enteritidis PT4 strain P125109 transposon mutant library consisted of about 140,000 insertion mutants (Li et al., 2020), and the *S.* Newport C4.2 transposon mutant library consisted of about 80,000 insertion mutants (de Moraes et al., 2018).

### 2.3 Comparative growth analysis

Growth curves on RTE muskmelon were assessed for all three barcoded transposon libraries (eight replicates each) and compared to their respective wild-type strains (five replicates each) at 8°C and 22°C to ensure that overall growth dynamics were not significantly altered in the mutant pools during the interaction with RTE muskmelon. Inoculation and sampling were performed as described for the screenings below. The growth data was subjected to an ANOVA test, where an adjusted *p*-value below 0.05 was considered significant.

### 2.4 Library screening on RTE muskmelon

For screenings on RTE muskmelon, 300 µl frozen stocks of the *S. enterica* barcoded mutant libraries were thawed and grown in LB^kan^ (Carl Roth; 60 μg/ml) at 37°C and 200 rpm until OD_600_ 1.0 was reached. One ml of this culture was diluted 1:10 in phosphate-buffered saline (PBS; Carl Roth) to reach approximately 8.0 log_10_ CFU/ml, centrifuged at 4.500 rcf for 5 min, the supernatant discarded, and the pellet resuspended in 5 ml PBS to create the inoculum.

Galia muskmelons (*Cucumis melo* L. *reticulatus*) were purchased at local grocery stores. For inoculation, muskmelon cubes were cut with sterile knives, and 10 g were weighed into sterile filter Stomacher^®^ bags (0.4 L, Avantor VWR International GmbH, Darmstadt, Germany). *No matrix* samples containing only the inoculum without the muskmelon were included as negative controls. RTE muskmelon samples and *no matrix* samples were inoculated with 5 ml of the inoculum to reach an initial concentration of approximately 7.0 log_10_ CFU/g or ml to ensure that the complexity of the library was maintained. The inoculum was evenly distributed over the muskmelon samples by manually massaging the bags, ensuring the muskmelon cubes were completely covered. Samples were incubated at 22°C and 8°C for 24 h and 5 days, respectively. No macroscopic changes were observed in the appearance of the muskmelon cubes during the incubation period.

Additionally, uninoculated RTE muskmelon samples were examined for total mesophilic aerobic bacteria (MAB; EN ISO 4833-2:2013) and *Enterobacteriaceae* (EN ISO 21528-2:2017; anaerobic incubation: O_2_ <0.1%, CO_2_ 7.0-15.0%) at every sampling time point. The absence of pre-existing *Salmonella* spp. in the purchased melons was confirmed by qualitative microbiological analysis according to EN ISO 6579-1:2017, A1:2020.

Mutant libraries were recovered after 1 h (t_1_), 7 h (t_7_) and 24 h (t_24_) for experiments at 22°C, and after 1 h (d_1_), 48 h (d_3_) and 96 h (d_5_) at 8°C. For this, the full muskmelon and *no matrix* samples were individually transferred into 90 ml LB^kan^ (Carl Roth) in 100 ml-Erlenmeyer flasks and incubated at 37°C and 200 rpm for 30 min to detach *Salmonella* from the matrix. After this, *Salmonella* populations were quantitatively assessed by preparing serial dilutions in PBS and plating on XLT-4 agar (Oxoid) that was then incubated at 37°C for 24 h. The RTE muskmelon samples in 90 ml LB^kan^ were then centrifuged at 4.500 rcf for 5 min to eliminate muskmelon juice residues, and the pellet was resuspended in 90 ml LB^kan^. Samples were then incubated for further 7.5 h at 37°C and 200 rpm before genomic DNA extraction for subsequent sequencing. Five biological replicates were performed.

### 2.5 Sequencing library preparation and data analysis

Aliquots of 40 µl from each sample were washed three times in sterile water, pelleted and then lysed with proteinase K (Sigma-Aldrich; 100 µg/ml) in 20 µl 10 mM Tris [pH 8.0] with 1 mM EDTA (Carl Roth) and 0.1% Triton X-100 (Carl Roth) for 2 h at 55°C. After enzyme inactivation at 95°C for 10 min, 5 µl were subjected to PCR with indexing primers targeted to the regions directly adjacent to the barcode on the transposon (0.4 µM Illumina-barcoded Primers L and V, listed in **Table S1**). The PCR was carried out using the following thermal cycling conditions: an initial denaturation step at 98°C for 2 min, followed by 10 cycles of denaturation at 98°C for 10 seconds, annealing at 65°C for 10 seconds, and extension at 72°C for 20 seconds. This was followed by additional 20 cycles consisting of denaturation at 98°C for 10 seconds and extension at 72°C for 20 seconds. A final extension was performed at 72°C for 2 minutes. The reaction was subsequently held at 20°C. The Q5™ Hot Start High-Fidelity 2x Master Mix (Item number M0494L; New England Biolabs, Frankfurt am Main, Germany) was added at a final concentration of 1x. Equal volumes of PCR products were pooled and purified using the QIAquick PCR Purification Kit (Qiagen, Item number 28106), following the manufacturer’s instructions. Illumina sequencing was performed as a dual-indexed paired-end 50, 100 or 150-base run on a NovaSeq 6000 with at least one million reads/sample.

Raw reads were binned based on their indexes and analyzed using custom Python scripts to identify and enumerate barcodes bordered by the expected invariable DNA sequence. Barcode insertions were compiled into an aggregated count for each disrupted gene. The differences in the aggregated abundances of the different mutants between time points, as well as between the matrix and *no matrix* samples for each sampling time point, were statistically analyzed using DESeq2 (Love et al., 2014) and log_2_-fold changes (fc) were reported (**Table S2**).

Mutations were considered to have a significant fitness effect for *S. enterica* on muskmelon if they met two criteria at the same sampling time point: (i) a log_2_-fc >|1.0| with a *p*_adj_*-*value ≤0.01 when comparing the growth on muskmelon to the growth in PBS (*no matrix* sample), and (ii) the same statistical threshold (log_2_-fc >|1.0|, adjusted *p*_adj_≤0.01) when comparing the inoculum to the muskmelon sample at the same sampling time point. For mutations identified at the final sampling timepoint (t_24_ or d_5_), criterion (ii) was also considered fulfilled if the comparison was made between the first and final muskmelon time points (t_1_ vs. t_24_ or d_1_ vs. d_5_) rather than the inoculum, thereby reflecting interaction mechanisms that emerged *after* the adaptation to the muskmelon environment. This stringent dual filtering was designed to identify genes with the most relevant roles for the interaction of the tested *Salmonella* serovars with muskmelons.

To generate gene lists for the KEGG (Kanehisa et al., 2023)- and GO (Ashburner et al., 2000; Thomas et al., 2022; Aleksander et al., 2023) analysis input, the above *p-*value criterion of *p*_adj_≤0.01 was relaxed to *p*_adj_<0.05. GO enrichment analysis was carried out using TopGO (Alexa and Rahnenfuhrer, 2024) and KEGG enrichment analysis used enrichKEGG from ClusterProfiler (Wu et al., 2021) in R (v.4.4.1). For KEGG results, pathways with *q-*values≤0.01 were considered significant. Resulting GO terms with classic Fisher *p*-values≤0.01 were considered significant.

### 2.6 Competition assays

To confirm the phenotype of candidate genes identified using TIS, single-gene deletion (SGD) mutants in *S*. Typhimurium 14028s were used in direct competition with the WT strain. These mutants were obtained from an existing SGD library(Porwollik et al., 2014) in STM that includes two mutants for a majority of all genes, one mutant harboring a kanamycin resistance gene in the sense direction of the deleted gene (SGD-Kan^R^) and one mutant harboring a chloramphenicol resistance gene in the antisense direction (SGD-Cm^R^). The WT (*S.* Typhimurium 14028s) used in the competition experiments was isogenic except for a tetracycline resistance cassette inserted in the neutral *malXY* genome location (WT_STM_-Tet^R^).

SGD mutants and the WT_STM_-Tet^R^ were grown in 5 ml LB^kan^, LB^cm^, and LB^tet^, respectively, at 37°C with 200 rpm shaking until approximately OD_600_ 0.9 was reached. For each selected gene, the different SGD cultures were mixed in a 1:1:1 ratio (or 1:1 if only one SGD mutant was available in the collection). The SGD-WT_STM_-Tet^R^ mix was then pelleted, resuspended in PBS, and 5 ml were inoculated onto 10 g RTE muskmelon at a concentration of approximately 7.0 log_10_ CFU/g and screened as previously described. Post-screening, samples were plated on LB^kan^, LB^cm^, and LB^tet^ agar and incubated at 37°C for 24 h to assess changes in population ratios. All mutants were similarly tested in competition assays in LB broth to investigate whether any genes played a role specific to growth on muskmelon or whether their role was universal in a nutrient-rich environment. Furthermore, some mutants were also tested in competition assays in PBS to confirm lack of differential death in the absence of nutrients. Competition assays in PBS and LB were inoculated at approximately 7.0 log_10_ CFU/ml. All cultures were tested for cross-resistance before being combined into a mixed inoculum.

The mutant phenotype was confirmed by comparing the changes in colony counts between samplings in the respective sense and antisense mutants and the WT_STM_-Tet^R^. If the competitive index (the ratio of mutant/WT ratios of timepoint/inoculum), CI, To confirm a negative fitness effect, both SGD mutants were required to display a competition index where CI+STDEV<1. In contrast, for a confirmed positive fitness effect, both mutants needed to adhere to the CI– STDEV>1 threshold. If only one SGD mutant was available, an expected single mutant behavior as outlined above was sufficient to confirm the phenotype observed in the transposon assay.plus standard deviation from at least three experiments remained below 1, a negative fitness effect was deemed confirmed. If the competitive index minus standard deviation remained higher than 1, a positive fitness effect was deemed confirmed. Competition assays were performed in triplicate, except when no fitness effect was observed in the first two replicates.

## 3. Results

### 3.1 Growth data

The barcoded transposon libraries grew 1.0 to 2.0 log_10_ CFU/g on muskmelon, while negative controls in PBS remained stable at 22°C and decreased slightly at 8°C **(Figure S1)**. The SEN mutant library showed reduced growth at 8°C compared to SNP and STM. In comparison with the WT, the transposon mutant libraries exhibited comparable growth on RTE muskmelons at 8°C and 22°C (**Figure S2**), except for the SNP library, which showed a statistically significant minor reduction compared to its corresponding WT (−0.3 log_10_ CFU/g, *p*-adj≤0.01) at d_1_, 8°C.

Mesophilic aerobic bacteria (MAB) and Enterobacteria on *Salmonella*-free RTE muskmelon increased by approximately 4.0 log_10_ CFU/g at 8°C and 5.0-6.0 log_10_ CFU/g at 22°C, with high variability among samples (**Figure S3**).

### 3.2 Transposon Insertion Sequencing data analysis

TIS screening of STM, SEN, and SNP libraries on RTE muskmelon identified functionally grouped mutants with significant fitness effects. All fitness data of the transposon assay after DESeq2 analysis are presented in **Table S2**, while the subset of genes with a significant fitness effect, as detailed in Materials and Methods, are displayed in **Table 2**. Exemplary volcano plots for inoculum/end of storage comparisons for the three *S. enterica* serovars are shown in **Figure 1** while the mutants’ dynamics across all analyzed time points at both temperatures are fully represented in **Figure S4**. These figures support the interpretation of temporal fitness patterns and provide a comprehensive overview of the log₂ fold changes of each significant gene and their associated significance for each comparison.

**Figure 1.**
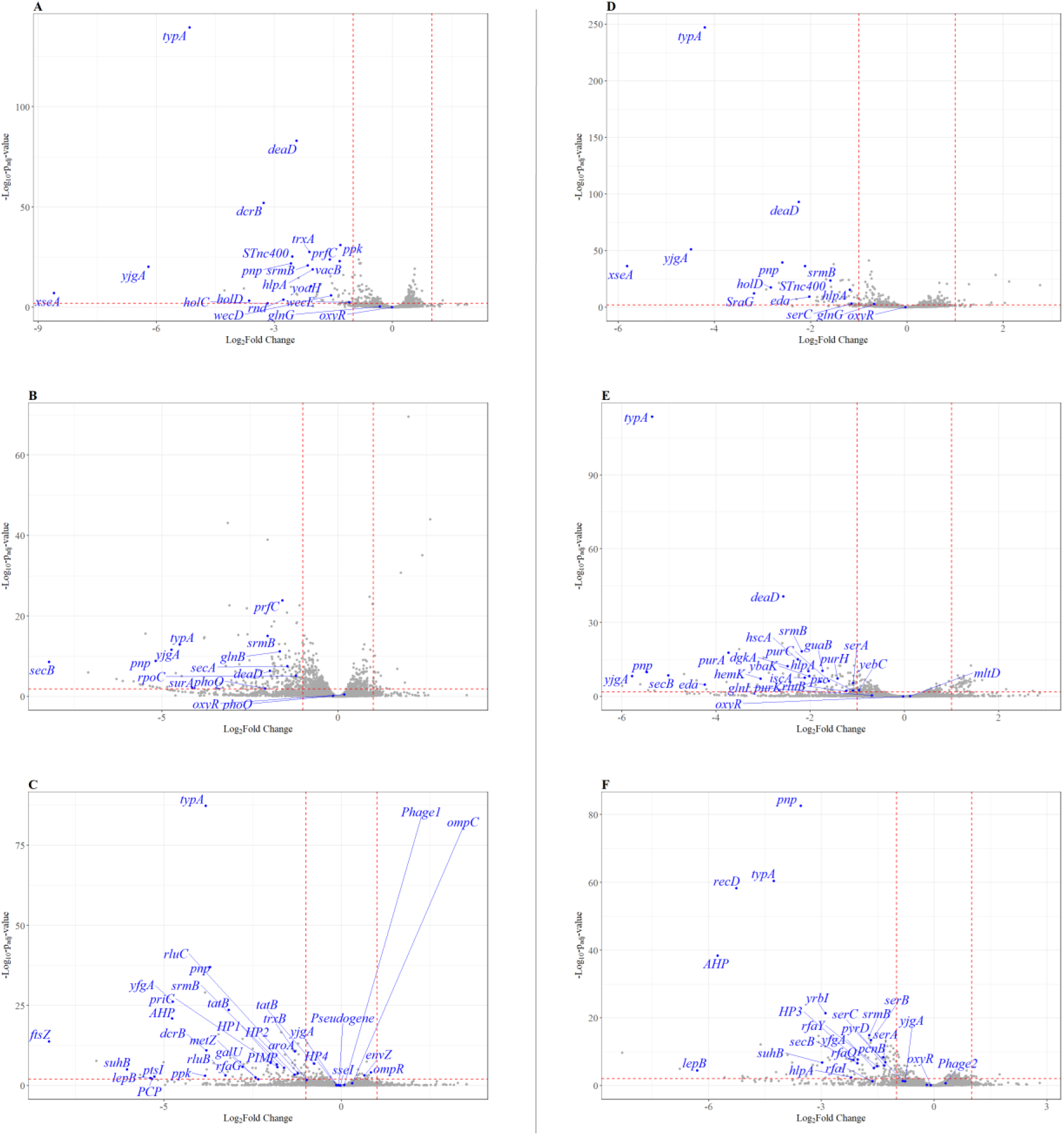
Changes in mutant abundances during the interaction of the *S. enterica* transposon mutant libraries on RTE muskmelon. Volcano plots depict genome-wide fitness effects by showing log_2_ fold changes in relative mutant abundance on the X-axis and corresponding adjusted *p*-values on the Y-axis when comparing muskmelon sampling time points. **Panels A-C** display results after 24 h at 22°C for *S.* Typhimurium 14028s (**A**), *S.* Enteritidis PT4 strain P125109 (**B**), and *S.* Newport C4.2 (**C**). **Panels D-F** present the same comparison after 4 days at 8°C for the respective strains: *S.* Typhimurium 14028s (**D**), *S.* Enteritidis PT4 strain P125109 (**E**), and *S.* Newport C4.2 (**F**). Each dot represents the aggregate mutant abundances for one specific gene. Mutants with significantly increased or decreased abundances are labelled in blue. Intergenic regions (IRs) with significant fitness effects were excluded. Additional time points and dynamics are shown in **Supplementary Figure S4**.

**Table 2.**
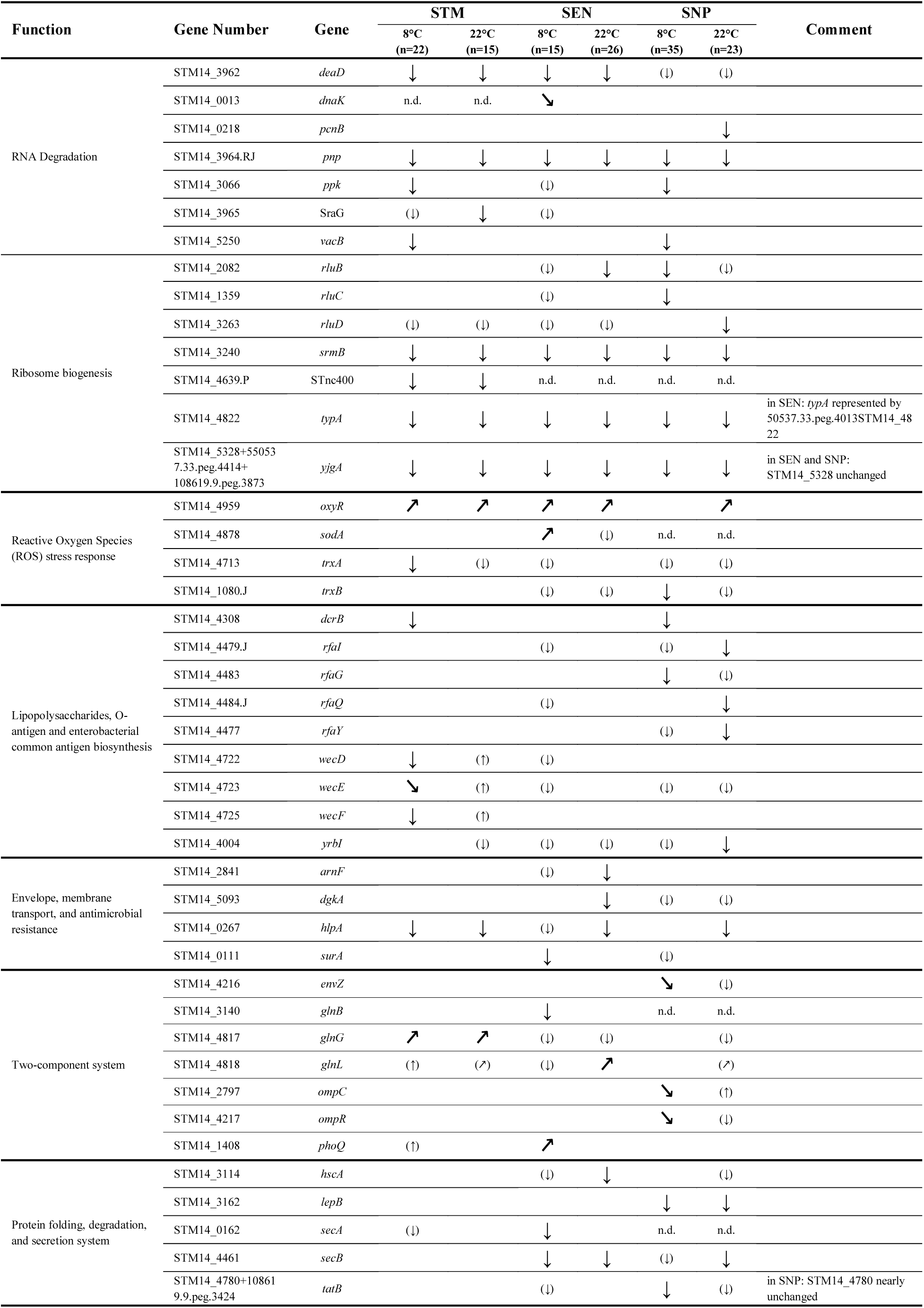
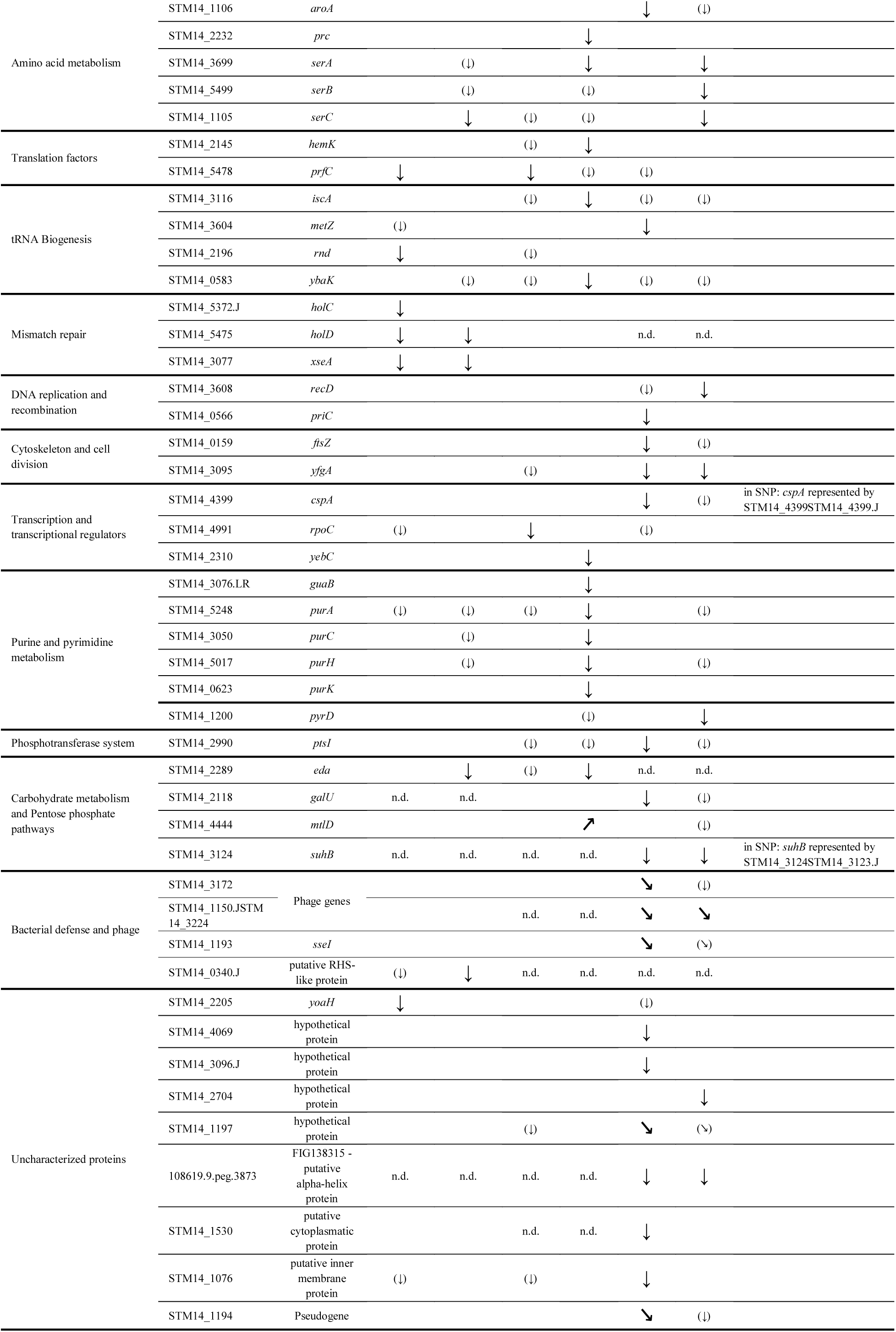
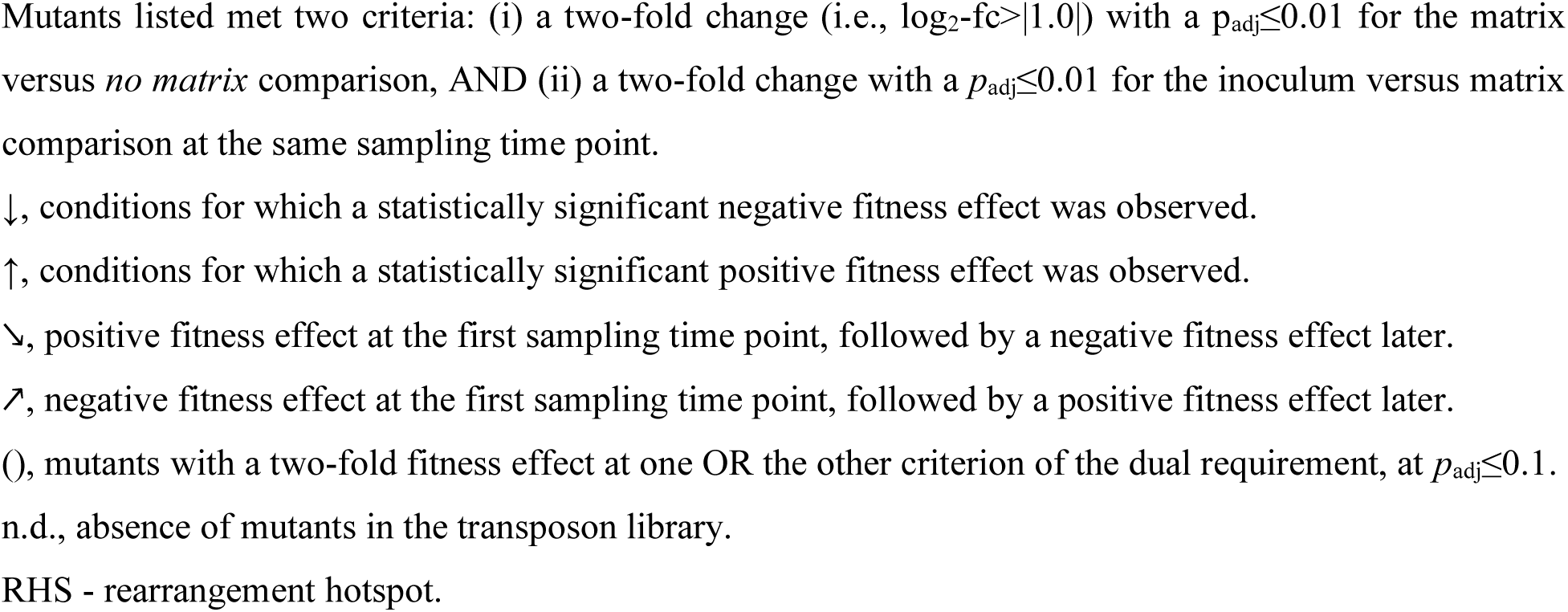
Genes with a significant fitness effect in *Salmonella enterica* serovars when grown on muskmelon, classified by function.

Three core genes (*pnp, typA,* and *srmB*) contributed to fitness in all three *S. enterica* serovars at both temperatures, while *hlpA* and *oxyR* mutants were found to be underrepresented in all three serovars at 22°C. For STM, mutants in 26 genes showed a significant fitness effect on RTE muskmelon, including 22 genes at 8°C and 15 genes at 22°C, with 11 overlapping. Most mutations had a negative fitness effect, except for *oxyR* and *glnG,* that switched from early negative to late positive fitness effects, and *wecE,* which showed the opposite trend. In SEN, mutants in 34 genes exhibited a significant fitness effect on RTE muskmelon. At 8°C, 15 genes were identified, while at 22°C, mutants in 26 genes showed fitness effects. Mutants in seven of these genes were significantly affected in both temperatures (*pnp, deaD, yjgA, srmB, typA, secB, oxyR)*. In SNP, mutations in 23 genes affected fitness on RTE muskmelon at 22°C, while 35 genes were implicated at 8°C. Temperature-independent fitness effects were observed for nine of these genes (*pnp, srmB, typA, yfgA, yjgA, suhB, lepB,* FIG138315 and one phage gene; **Table 2**).

In all three serovars, mutants in RNA degradation genes, ribosome biogenesis, envelope and membrane transport, and tRNA biosynthesis showed a statistically significant difference after growth on muskmelon. For some categories displayed in **Table 2**, statistical significance, as defined in our criteria, was only observed for genes in one or two serovars. As an example, mutants in genes of the purine and pyrimidine pathways were statistically significantly affected in growth in SEN, but not in STM or SNP, and cytoskeleton and cell division gene mutants cleared the significance threshold in SNP but not in the other two serovars (**Table 2**). However, the number of transposon mutants in each gene and their abundance in the library differ among strains, so these differences should not be misinterpreted as confirmation of a difference in the role of that pathway between strains.

We performed KEGG enrichment analyses to gain a better insight into pathways significant for the growth of *Salmonella* on muskmelon. Significantly enriched KEGG pathways (*q*≤0.01) and associated GO terms (*p*≤0.01) are summarized for all three serovars in **Table 3**, with full results in **Table S3.**

**Table 3.**
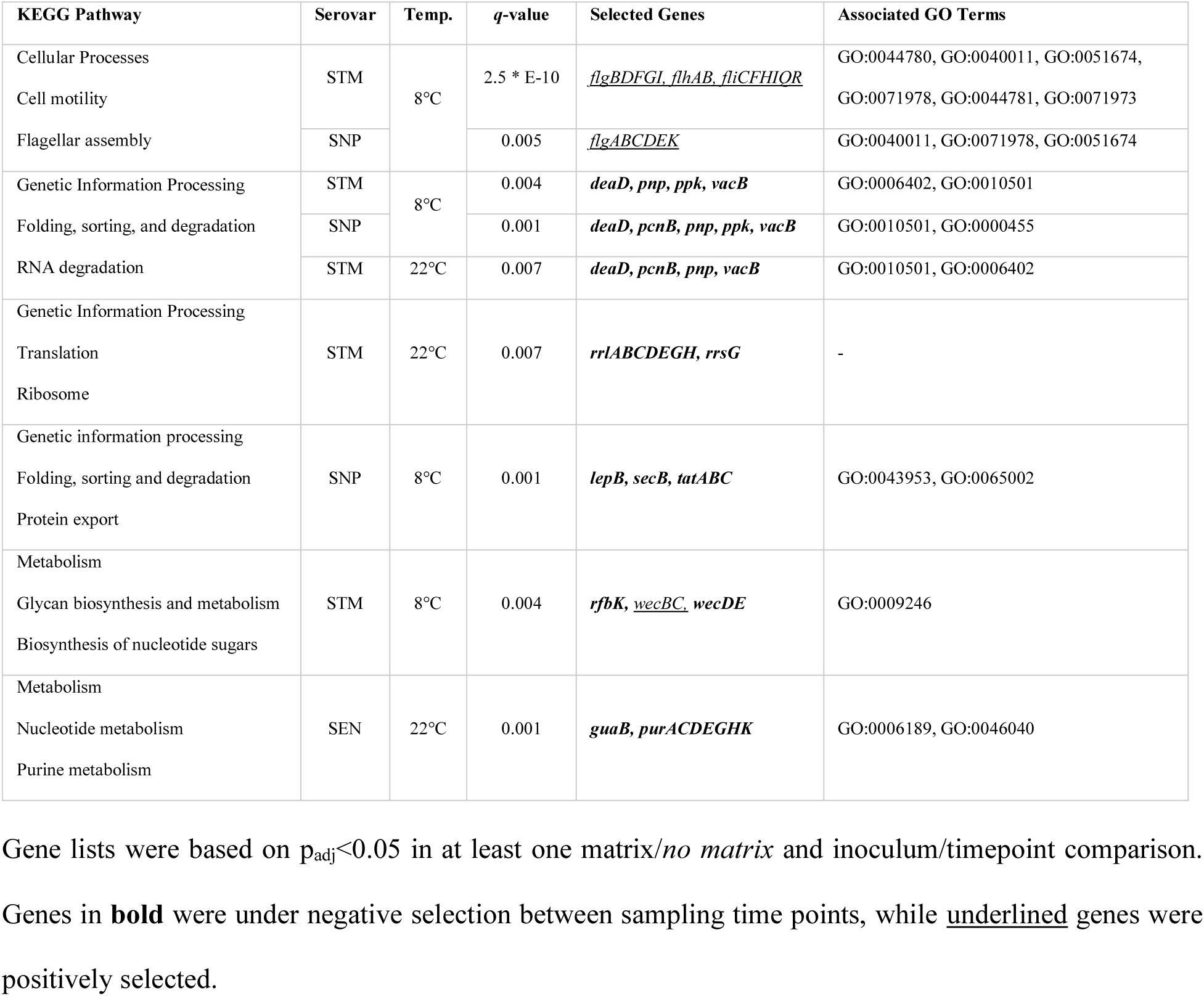
Significantly enriched KEGG pathways and associated GO terms for *S.* Typhimurium 14028, *S.* Enteritidis P125109, and *S.* Newport C4.2 on RTE muskmelon.

### 3.3 Competition assays confirm genes with a fitness effect

A total of 25 genes with a fitness effect in STM, SEN, or SNP on RTE muskmelon were selected from the *S.* Typhimurium 14028 SGD mutant collection (Porwollik et al., 2014) for an extensive set of competition assays (**Table 4**). Four of these genes (*serA*, *serB, ybaK*, and *purA*) displayed changes in STM that were not as significant as in SEN or SNP but were tested to verify whether the results of the transposon assays were, in fact, a representation of the true diversity between *Salmonella* serovars. Two genes that exhibited temporal abundance changes in STM, *oxyR* and *glnG*, were also included in these competition assays, along with *rfbI_2*, which showed no significant changes in any of the three strains analyzed. A final gene, *wecF [D]*, showed opposite effects in the two different incubation temperatures in STM. However, significance (*p*_adj_<0.01) was only achieved at 8°C, while at 22°C only the inoculum/t_24_ comparison was significant.

**Table 4.**
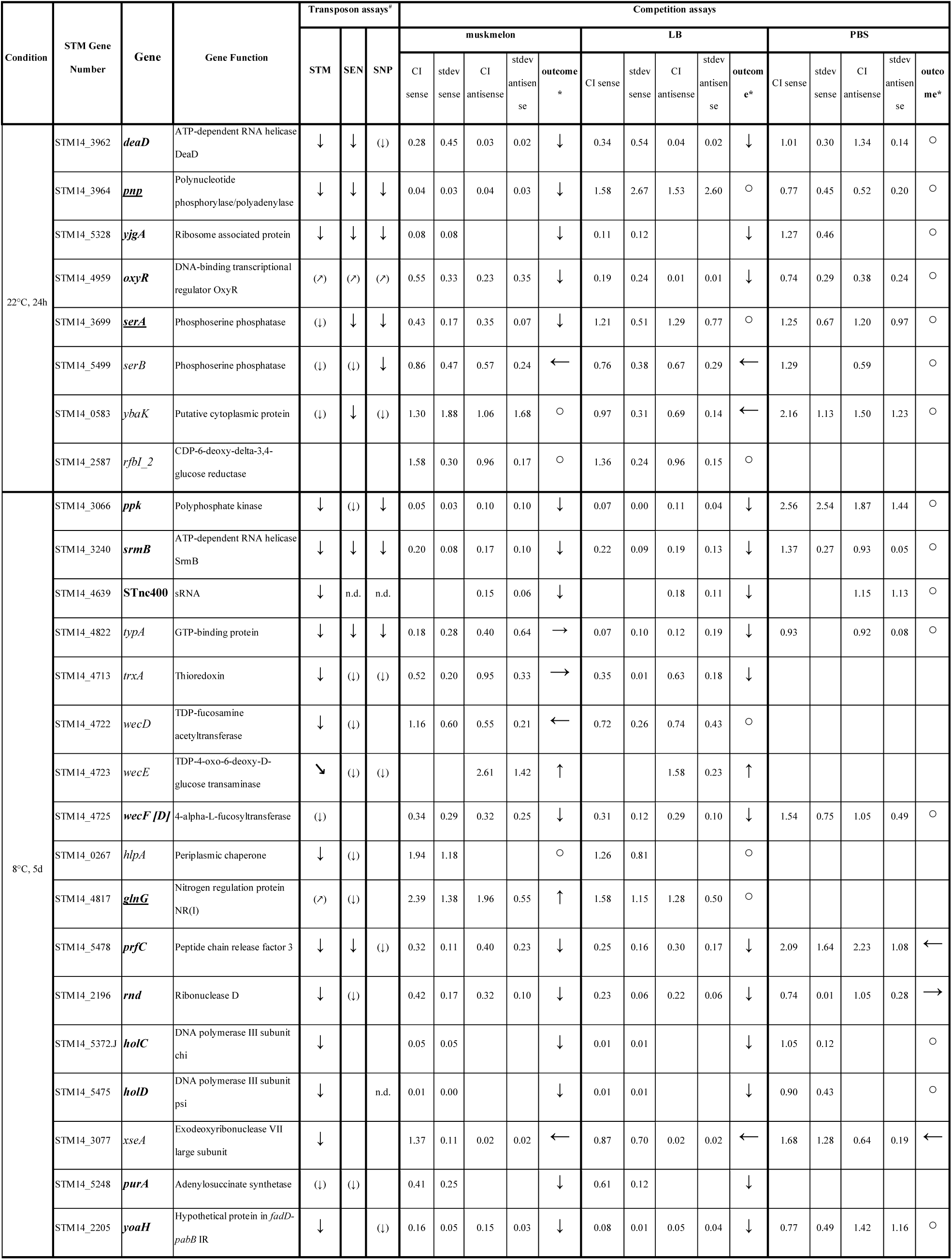
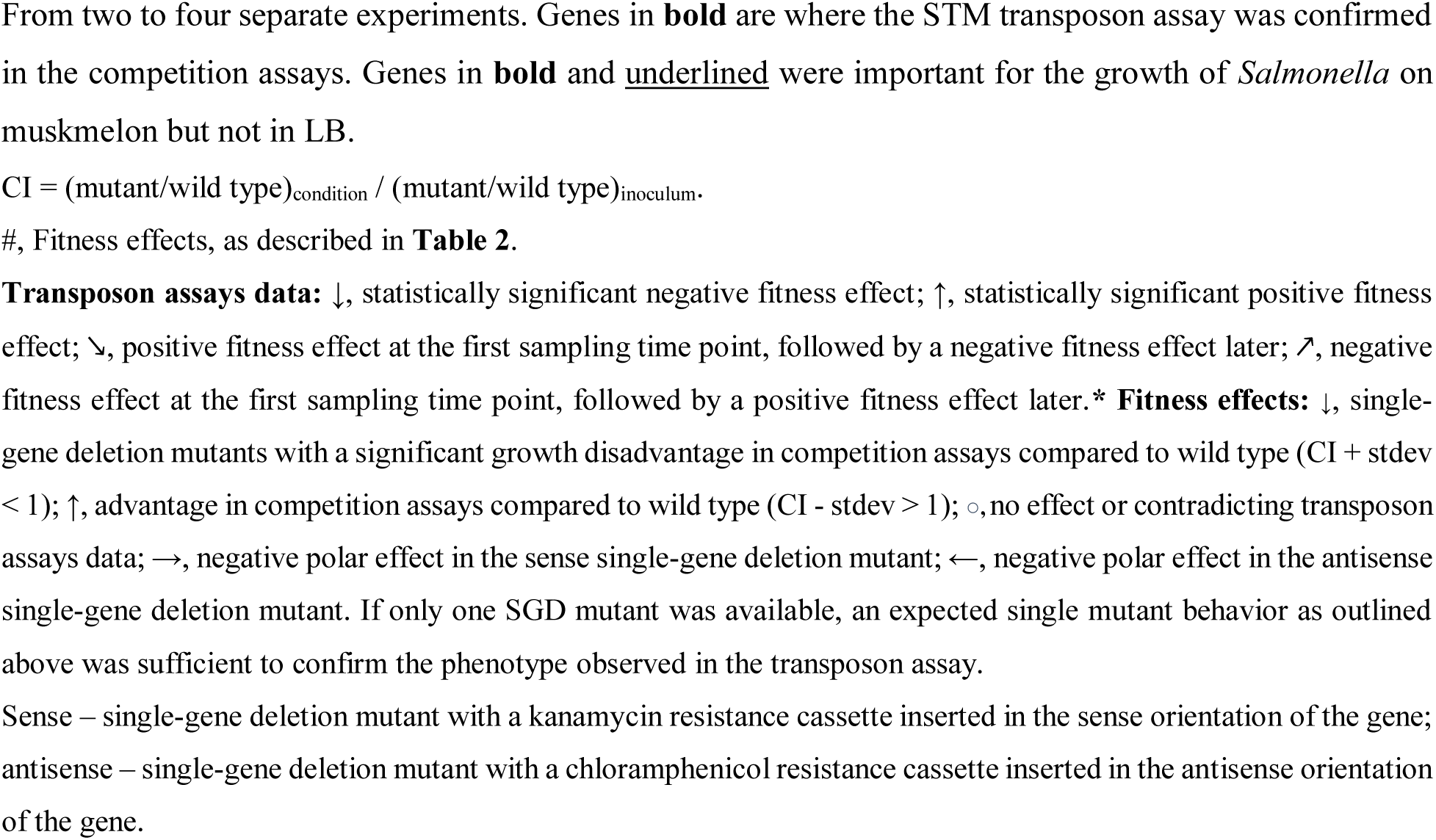
Fitness outcomes of selected *S.* Typhimurium 14028s single-gene deletion mutants under competition against the wild type.

For 18 of these 25 genes, we had two single-gene deletion mutants available for each gene, one where a kanamycin resistance (Kan^R^) cassette replaced the gene in the sense orientation and one where a chloramphenicol cassette (Cm^R^) replaced the gene in the antisense orientation. For five genes (*yjgA, hlpA, holC*, *purA,* and *holD*), we only had a Kan^R^ mutant available, while for *wecE*, our collection only harbored a Cm^R^ mutant. Competition assays were performed at the temperature where transposon data suggested a stronger fitness effect. Assays were performed on muskmelon and rich LB media, to verify whether any fitness effects occurred in any rich nutrient environment or were potentially specific to growth on muskmelon. For selected genes, the fitness effect observed in muskmelon or LB was also tested in PBS, where the nutrient-restricted conditions only confer a growth difference due to differential death. The results are summarized in **Table 4**.

As shown in **Table 4**, the observed fitness effects in the transposon assays were confirmed for 15/24 genes. Notably, *serA* and *pnp* deletions showed a negative fitness effect during the competition with the WT on muskmelon that was not detected in LB. Similarly, the positive fitness effect of transposon insertions in *glnG*, observed in STM at 8°C on d_5_, was confirmed in competition assays on muskmelon and not on LB, although with high standard deviations. Further replicates would be needed to illuminate this difference further. Mutants in *xseA* exhibited a strong polar effect in LB and RTE muskmelon, where one single-gene deletion mutant confirmed the expected debilitated phenotype on muskmelon, while the other suggested a growth advantage. This may be due to the different orientations of the constitutive promoter introduced into the gene during the generation of the two mutant constructs. For four genes (*wecD, trxA, typA*, and *serB*), one SGD mutant confirmed the expected phenotype while the other showed no significant change.

## 4. Discussion

This study investigated *S. enterica* fitness determinants on RTE muskmelon at room temperature and during abusive cold storage at 8°C using TIS in three serovars: *S.* Typhimurium ATCC 14028, *S.* Enteritidis PT4 strain P125109, and *S.* Newport C4.2. The growth potential was comparable to previous studies with *S. enterica* on muskmelons (Ukuku and Sapers, 2007; Huang et al., 2015; Feng et al., 2017), with minor variations probably related to strain differences, processing conditions, and time-temperature combinations. The slight reduction in SNP growth at d_1_ (8°C) may be due to phage lysis under stress, as phage genes were negatively selected at this temperature.

Limited data is available on the background microbiota in muskmelons and their growth during storage. Compared to our data, higher initial MAB counts (3.5-4.0 log_10_ CFU/g total aerobic bacteria compared to ∼2.0 log_10_ CFU/g in our study) and coliforms (slightly below 2.0 log_10_ CFU/g compared to 0.5-1.0 log_10_ CFU/g *Enterobacteriaceae* in our study) were reported in previous studies (Ukuku and Sapers, 2007; Song et al., 2024). This is likely attributable to the handling of muskmelon samples, which were cut into cubes under sterile conditions.

TIS revealed shared fitness determinants across all three serovars, particularly genes involved in RNA metabolism, ribosome and amino acid biogenesis, oxidative stress response, and cell integrity and membrane transport. Serovar-specific fitness effects were observed for mutations in genes related to protein folding and transport, purine and pyrimidine metabolism, DNA replication, recombination, and mismatch repair. Although these differences have not been further analyzed and may be due to differences in the number of transposon mutants in each gene and their abundance in the library, this approach allowed us to spread a wider net, capturing genes under selection that would have been missed with only one TIS library. Future work may analyze single-gene mutant orthologs to elucidate whether candidate differences between serovars are indeed differences.

Mutations in genes involved in **RNA degradation** (*pnp*, *deaD, vacB,* SraG*, ppk, dnaK, pcnB)* **and ribosomal homeostasis** (*typA, srmB, yigA, rluBCD, STnc400*) had significant fitness effects across serovars.

PNPase (exoribonuclease polynucleotide phosphorylase (*pnp*)), a conserved RNA degradosome component, regulates mRNA degradation, tRNA processing, and small RNA turnover (Clements et al., 2002). PNPase is critical for bacterial cold-shock responses, although *S. enterica pnp* mutants exhibit milder cold sensitivity, consistent with its temperature-independent fitness effect on muskmelon. Its relevant role during the interaction on muskmelon was confirmed with a muskmelon-specific negative fitness effect absent in nutrient-rich media in the competition experiments against the WT.

DeaD and SrmB are DEAD-box RNA helicases (Charollais et al., 2003; Vakulskas et al., 2014). Both *srmB* and *deaD* mutants exhibit cold-sensitive growth defects in *E. coli* (Charollais et al., 2003)*, Pseudomonas syringae* (Hussain and Ray, 2024), and *Yersinia pseudotuberculosis* (Jiang et al., 2019), consistent with our findings, although the effect was observed at both temperatures. Beyond cold stress, DeaD and SrmB participate in post-transcriptional regulation, including the carbon storage regulatory system in *E. coli* (Vakulskas et al., 2014), which may explain their temperature-independent fitness effects. Co-transcription of *deaD* and *pnp* occurs in certain bacterial species but is absent in others (Jones et al., 1996; Favaro and Dehò, 2003). This is mirrored in our data, where *deaD* disruption affected fitness only in STM and SEN but not SNP. This phenotype was confirmed on both muskmelon and nutrient-rich media in the competition experiments, and is therefore not melon-specific.

PPK (Polyphosphate kinase) is highly conserved across bacteria. It catalyzes inorganic polyphosphate (polyP) synthesis, regulates RNA degradation and mRNA biogenesis (Zhu et al., 2005), and contributes to *S. enterica* growth, survival, and virulence (McMeechan et al., 2007). PolyP, a high-energy reservoir, plays a key role in STM and SNP muskmelon interactions but shows no fitness effect in SEN. Serovar-specific differences in *ppk* function were previously observed (McMeechan et al., 2007), with *ppk* mutants exhibiting a greater growth defect in *S.* Typhimurium than *S.* Gallinarum, suggesting variations in energy homeostasis or alternative energy use. PolyP synthesis occurs mainly in stationary phase (McMeechan et al., 2007), likely balancing metabolism by regenerating ATP through the removal of inhibitory polyP and nucleotide diphosphates (Blum et al., 1997). On muskmelon, *ppk* mutants displayed a fitness effect only at 8°C and later time points, indicating polyP accumulation is critical under prolonged cold stress, possibly to maintain mRNA degradation homeostasis, a phenotype confirmed for muskmelon and nutrient-rich media. Nevertheless, polyP is also known to be essential for *S. enterica* long-term survival in LB, even at 37°C (Kim et al., 2002). Additionally, *vacB* mutants (exoribonuclease R), deficient in RNA degradation and processing as well as in ribosome biogenesis (Hammarlöf et al., 2015), exhibited a fitness effect under the same conditions as Δ*ppk* mutants. Taken together, these findings indicate the importance of RNA and ribosomal homeostasis in *S. enterica* environmental adaptation.

In the presence of **reactive oxygen species (ROS)**, OxyR acts as key regulator of oxidative stress. Δ*oxyR* mutants exhibited a dynamic fitness effect that was initially strongly negative but shifted to a positive selection over time in SEN and STM at both temperatures and in SNP at 22°C.

This pattern suggests *oxyR* disruption initially impairs bacterial adaptation but later confers a selective advantage. Members of the *oxyR-*regulon (e.g., *trxAB*)(Prieto-Alamo et al., 2000) also showed a negative fitness effect in STM and SNP at 8°C, reinforcing the role of oxidative stress responses in muskmelon adaptation. The competition assays mirrored the transposon results, albeit with high variability, further emphasizing the complex environmental regulation of *oxyR*. While a negative fitness effect was also observed in LB, this effect could be due to the H₂O₂ content of the medium. While plants generally generate ROS as a defense response (Kuźniak and Kopczewski, 2020), cut muskmelon pieces are unlikely to retain this response. Thus, the *oxyR*-effect may relate to alternative metabolic adaptations (Cota et al., 2012; Cota et al., 2015) that shall be further investigated in future studies.

**Cell wall integrity** and **membrane transport** components were also identified. Several genes involved in LPS and membrane biosynthesis (*rfaG, dcrB*) and enterobacterial common antigen (ECA) biosynthesis (*rfaG, wecDEF, dcrB*) exhibited fitness effects, particularly at 8°C, although for SNP, the negative fitness effect was prevalent at 22°C. While the fitness effect at 8°C may represent an adaptation to altered membrane fluidity at cold temperatures (Ricke et al., 2018), a special need for the membrane LPS or specifically for the ECA in SNP during its interaction with muskmelon at room temperature may be of importance. LPS biosynthesis was shown to be important for *S. enterica* survival on pistachios (Jayeola et al., 2020), tomatoes (de Moraes et al., 2017), and mouse colonization (de Moraes et al., 2017), reinforcing their broad role in environmental adaptation.

Disruptions in membrane transport (*secAB, lepB, hlpA, hscA, surA, tatB*) impaired bacterial fitness on muskmelon, indicating that efficient protein translocation is critical for survival. The chaperone SecB, required for protein targeting to the SecYEG transport system, was particularly important in SEN and SNP. The outer membrane protein HlpA also exhibited a fitness effect at 22°C, although competition assays did not confirm this phenotype. Nevertheless, the enrichment of membrane transport genes suggests that maintaining protein export balance is essential; either the accumulation of metabolites in the cytoplasm or the depletion of functional proteins in the periplasm may be detrimental for *S. enterica* growth on muskmelons.

**Two-component systems (TCSs)** and **motility** genes were also overrepresented in the TIS data. Mutants in various TCSs showed fitness effects at 8°C, suggesting a role under cold stress. Nitrogen assimilation TCS mutants (*glnG* in STM, *glnBL* in SEN) showed significant fitness effects, while *phoQ* mutants (sensing antimicrobial substances and Mg²⁺ starvation) appeared to be affected only in SEN. In SNP, osmolarity regulators (*envZ, ompR, ompC*) exhibited fitness effects. Positive selection for *glnG, phoQ, envZ*, and *ompR* mutants during part of the interaction period suggests a fitness advantage.

Positive selection was also observed for mutations in flagellar assembly genes, likely aiding adaptation to the plant environment (Sanguankiattichai et al., 2022). This correlates with the results of a previous study, where *S.* Typhimurium 14028 largely downregulated flagellin production, while a subset of cells exhibited high flagellin expression inside the plant (Zarkani et al., 2020).

**Protein synthesis** was relevant for the interaction of *S. enterica* with muskmelons, as inactivation of genes involved in the serine (*serABC, prc)*, aromatic amino acids (*aroA),* and glutamine metabolism (*glnB)* resulted in serovar- and temperature-dependent fitness effects. Additionally, genes related to tRNA (*rnd, iscA, ybaK, metZ*) and translational regulation (*prfC, hemK*) were also identified. Both *serA* and *glnB* were previously identified as necessary for tomato colonization (de Moraes et al., 2017), suggesting a critical role for scavenging serine and glutamine from the food matrix. The positive fitness effect observed at later time points for *glnG* mutants, a gene coding for the nitrogen regulation protein NR(I), was confirmed in the competition assays to be muskmelon-specific, suggesting that a switch to alternative nutrient sources or alternative regulation mechanisms of the nitrogen metabolism may be beneficial for growth on muskmelon. Perhaps nitrogen assimilation is unnecessary for the fitness of *S. enterica* on muskmelons. Interestingly, competition experiments with individual *serB* mutants revealed a polar effect in the antisense single-gene deletion mutant, further highlighting *serA* as a key fitness determinant on muskmelon.

*ΔaroA* mutants have demonstrated a fitness effect in SNP and SEN under desiccation stress on low-moisture foods (Jayeola et al., 2020). Since *aroA* mutants only showed an effect at 8°C, a role in cold stress could be hypothesized, with a potential link to TypA as a tyrosine-based enzyme with a role in cold-shock response (Fan et al., 2015). A growth defect for *E. coli met-*mutants was previously demonstrated (Kenri et al., 1992; Tiefenbacher et al., 2024). While the *de novo* biosynthesis of amino acids was also found to be critical for the persistence of *S. enterica* on tomatoes, *S. enterica* appeared to prioritize amino acid scavenging during animal infections (de Moraes et al., 2017; de Moraes et al., 2018).

Mutants in genes with a role in **purine metabolism** were strongly repressed in SEN at 22°C. Notably, *ΔpurA* exhibited nearly four log_2_-fc reductions, suggesting *purA* may be essential for growth. The purine biosynthesis was previously implicated in *S.* Typhimurium colonization of tomatoes (de Moraes et al., 2017; de Moraes et al., 2018), and competition experiments confirmed a negative fitness effect for STM Δ*purA* mutants on ready-to-eat muskmelon and nutrient-rich media.

The *pyrD* transposon mutants, deficient for this gene involved in the *de novo* pyrimidine metabolism and linked to both purine and pyrimidine *de novo* synthesis (Wilson and Turnbough, 1990; Vial et al., 1993), exhibited a fitness effect in SNP. Pyrimidine metabolism is required by *S.* Typhimurium infection in animals (de Moraes et al., 2017; Yang et al., 2017), and proliferation in tomatoes (de Moraes et al., 2017). However, our study found fitness effects only in SNP, although associated GO terms were also enriched for SEN and SNP. This might suggest an ability to scavenge pyrimidines from the environment for certain *S. enterica* strains (de Moraes et al., 2018).

Mutants in genes related to **Fe-S-cluster biosynthesis** showed fitness defects in SEN and SNP. These clusters play essential roles in electron transfer, substrate binding, Fe-S storage, gene expression, and enzyme activity (Johnson et al., 2005). Disruption in *hscA* and *iscA*, involved in Fe-S cluster assembly (Yang et al., 2006; Vickery and Cupp-Vickery, 2007), as well as *aroA,* required for siderophore synthesis and iron uptake, were detrimental during muskmelon interaction. Similar iron uptake genes were previously linked to *S. enterica* proliferation in tomatoes (Nugent et al., 2015), and *iscA* was relevant during alfalfa sprouts’ colonization (Holden et al., 2024), suggesting a shared iron requirement for fresh produce colonization.

Our analyses during the competition assays revealed that the fitness effects observed during the interaction of several STM mutants with muskmelon were similar to those observed in LB medium, suggesting that muskmelon provides conditions resembling a rich nutrient medium for *S. enterica*.

Additionally, several polar effects were revealed. One example illustrating this dynamic is the characterization of *xseA,* which exhibited a strong temperature-independent reduction in STM mutants during the transposon screen. However, in the follow-up competition assays, the mutation revealed evidence of a polar effect of the single-gene deletion mutant with insertions in the antisense orientation reproducing the fitness disadvantage observed in the screen. An effect that was identical for muskmelon and nutrient-rich media, but much milder in PBS, suggesting this is a nutrient-dependent fitness effect. The observed phenotype may be attributable to the genes located upstream in the genome, *guaA* and *guaB,* involved in the purine biosynthesis pathway. While these genes demonstrated some degree of impact across different screening conditions, only *guaB* met the stringent criteria for a significant fitness effect in *S.* Enteritidis at 22 °C, highlighting the need for further targeted analyses of adjacent or co-regulated genes in future investigations. A polar effect in the sense single-gene deletion mutant of *typA* was observed. Downstream of this gene is a large operon comprising six genes encoding poorly characterized proteins, which may be the subject of further investigation in future studies. Another polar effect in the sense SGD mutant was observed for *trxA.* Although the *rho* operon is located downstream of this gene, they are not part of the same operon. Further analysis is needed to clarify the cause of this effect.

An important limitation of our study was the high inoculation loads required to maintain the complexity of the transposon library, which was mitigated to some extent by conducting direct competition assays between individual candidate mutants and the WT. Future studies could focus on individual genes, allowing lower inoculation levels and a more precise assessment of growth potential. A further limitation is that several genes identified in the transposon screen were either unavailable as single mutants in the collection (*serC, secAB, vacB)* or not confirmed in competition assays (*hlpA, serB, ybaK)*. The latter may be due to sensitivity differences between the two methods: TIS detects subtle fitness effects within a mutant library, whereas competition assays measure fitness relative to the WT, potentially overlooking small but biologically relevant differences. Moreover, hitchhiking mutations within the mutant pool may obscure individual gene contributions, as complete mutant purity cannot always be ensured.

## 5. Conclusions

This study identified key fitness determinants enabling *S. enterica* survival on muskmelon, revealing both conserved and serovar-specific adaptation mechanisms. RNA metabolism, *de novo* amino acid and ribosome biogenesis, oxidative stress response, and cell wall integrity emerged as central survival factors, with additional metabolic adaptations influencing fitness under specific conditions. Many of these mechanisms align with known plant-endophyte interaction pathways, including ROS detoxification, nitrogen fixation, flagella, and cell wall components such as LPS, surface-associated proteins, or transporters (Pinski et al., 2019). Future studies should investigate the metabolic interplay between *S. enterica* and plant-derived nutrients to further elucidate bacterial survival strategies in fresh-produce environments.

## Acknowledgements

We would like to express our gratitude to Verena Hohenester for her invaluable technical support during this study. Additionally, we extend our thanks to Barbara Fritz, Johanna Dietz, Erika Altgenug, René Mamet, and Larissa Klose for their contributions and assistance with the competition experiments.

## 6. Supporting Information

Supplementary Figure S1. Growth of the *Salmonella enterica* barcoded transposon mutant libraries (lib) and the corresponding wild types (wt) on ready-to-eat muskmelon at 22°C (A) and 8°C (B).

Supplementary Figure S2. Quantitative assessment of the *Salmonella enterica* barcoded transposon mutant libraries.

Supplementary Figure S3. Quantitative assessment of the background microbiota on the ready-to-eat muskmelon samples.

Supplementary Figure S4. Changes in the aggregated mutant abundances throughout the interaction period of the barcoded transposon mutant libraries on ready-to-eat muskmelon.

Supplementary Table S1. Primers employed for library preparation prior to Illumina sequencing.

Supplementary Table S2. DESeq2 statistical analysis for the transposon insertion sequencing experiments of the three *Salmonella enterica* libraries on muskmelon.

Supplementary Table S3. Results of the KEGG- and GO-enrichment analysis.

## 7. Declaration of generative AI and AI-assisted technologies in the writing process

During the preparation of this manuscript, the authors used ChatGPT-4o to enhance readability and clarity. All content generated with these tools was thoroughly reviewed and edited by the authors, who take full responsibility for the final version of the manuscript.

